# Personality-specific pathways from peer victimization to adolescent alcohol misuse: A multilevel longitudinal moderated mediation analysis

**DOI:** 10.1101/2021.04.16.440234

**Authors:** Flavie M. Laroque, Elroy Boers, Mohammad H. Afzali, Patricia J. Conrod

## Abstract

Peer victimization is common in adolescence and have been associated with a broad variety of psychopathology and alcohol use. The present study assessed whether peer victimization has a time-varying effect on alcohol use through internalizing and externalizing symptoms and whether this indirect association throughout time is moderated by personality. This 5-year longitudinal study (3,800 grade 7 adolescents) used Bayesian multilevel moderated mediation models: independent variable was peer victimization; moderators were four personality dimensions (anxiety sensitivity, hopelessness, impulsivity, and sensation seeking); internalizing symptoms (anxiety, depressive symptoms) and externalizing symptoms (conduct, hyperactivity problems) were the mediators; and alcohol use, the outcome. Results indicated significant between, within, and lagged effects on alcohol use through internalizing and externalizing symptoms. There was significant between and within effects on alcohol use through internalizing symptoms for adolescents with high anxiety sensitivity and hopelessness, and significant between, within, and lagged effects on alcohol use through externalizing for adolescents with high impulsivity and sensation seeking. These findings implicate two risk pathways that account for how peer victimization enhances alcohol use risk and emphasize the importance of personality profiles that can shape the immediate and long-term consequences of victimization.

## Introduction

### Peer victimization and alcohol use

Peer victimization is defined as being a target of repeated peer in physical, verbal, or psychological way by perpetrators who intend to cause harm, and is based on an imbalance of power (Olweus, 1993). Major public concerns have been raised about the impact of peer victimization on short-term and long-term mental health (Evans-Lacko et al., 2017; van Geel et al., 2014; Woo et al., 2019). One particular concern is the effect of peer victimization on adolescent alcohol use. Adolescence is a period of physical and psychological transition, during which adolescents are vulnerable to the effects of alcohol use, the most commonly used substance during this period of time (Johnston et al., 2018). Early use of alcohol is associated with physical injury and risky health behaviors, its harmful effect on health may also interfere with brain development (Hill et al., 2000). Because of their respective potential for harm, recommendations are made to reduce peer victimization and prevent the early onset of alcohol use during adolescence. The association between the two has been investigated; however, there are three noteworthy key gaps in the relevant literature: (1) the use of longitudinal and sensitive developmental designs that differentiate common vulnerability from concurrent and longitudinal relationships (overall, short-term; long-term) to clarify the nature of temporal precedence in these associations, (2) mechanisms by which long-term relationships are maintained (paths) and (3) specificity of the relationships. This study aimed to address these three gaps in the literature using an exceptional database that allows for sensitive analysis of time-dependent variations in victimization, mental health and substance use throughout the course of adolescence.

### Effect of peer victimization on alcohol use

All pertinent literature on the issue has been summarized in a systematic review from Maniglio et al. (2017) and a meta-analysis from Moore et al. (2017). The latter emphasized that previous studies did not successfully disentangle the association because of methodological discrepancies, but also because of poor quality design: small sample size, important confounders not taken into account, or no time-varying associations tested. Overall, the main constraint was the cross-sectional design for most studies (108 cross-sectional versus 57 prospective cohorts). These issues may threaten the accuracy, reliability, and validity of the results, leading to mixed evidence. Some studies reported that peer victimization is associated with a reduced risk of engaging in harmful alcohol use (Moore et al., 2014; Nansel et al., 2001), others found no associations (Kelly et al., 2015; Quinn, Fitzpatrick, Bussey, Hides, & Chan, 2016; Tomczyk, Hanewinkel, & Isensee, 2015), whereas others suggest that being bullied may result in an increased probability of later harmful alcohol use, even when important confounders such as use of drugs and peer drinking was controlled (Swahn et al., 2011; Tharp-Taylor et al., 2009). When pooled analysis was performed in the meta-analysis, a “*possible causal*” association between peer victimization and alcohol use was found (Continuous Update Project Expert Report, 2018). The most recent studies investigating direct associations remain cross-sectional: being the target of peer victimization, independent of adverse childhood experiences, was associated with increased odds of intoxication in the past 30 days compared to adolescents who did not experienced peer victimization (Afifi et al., 2020), and was associated with an increase in the adulthood prevalence of alcohol use disorders (Woo et al., 2019). Overall, these inconsistencies could be due to the cross-sectional nature of these studies and might also be partly due to confounding variables and individual differences that are not taken into account or investigated. Further longitudinal research that establish temporal inference is needed to better explore the underlying time-varying mechanisms (when and how) peer victimization is associated with alcohol use. Testing hypotheses of temporal dynamics requires certain design qualities: 1. *Temporal precedence* (e.g., time-varying relationships are demonstrated); 2. *Robust analytic methodology* (e.g., structural equation modeling involving random intercepts to control for common vulnerability over time) 3. *Mediation path* (how long-term relationships are maintained) and 4. *Specificity* (under which conditions long-term relationships are observed). The key challenge is to find the right model that will disentangle this complex maladaptive relationship. Thus, the current study aims to bring new evidence for time-varying associations by investigating overall, short-, and long-term associations between peer victimization and alcohol use through mediation and moderation processes.

### Psychopathology as a mediator

Integrating potential mediation effects is of particular interest for two reasons: (1) it provides important information for why and how relationships occur, (2) it allows to alleviate the underestimation of the effects sizes (Holbert & Stephenson, 2003). Broadly, it is the combination of direct and indirect effects that build the association between peer victimization and alcohol use. Interestingly, Moore et al. (2017) concluded that there was “*convincing evidence*” for a temporal relationship between prior peer victimization experiences and other psychopathological outcomes later, such as anxiety and depression, which are also known to further predict alcohol use (Hussong et al., 2011), indirectly implicating a mediation path from peer victimization to alcohol use. The existence of this pathway might also explain the inconsistencies between studies: it is possible that effects on alcohol use will emerge later after the effects of the mediating variable have accumulated over time. As suggested by the self-medication hypotheses (the internalizing symptoms pathway), on which most studies are based, some adolescents might use alcohol as a maladaptive attempt to cope with or escape negative emotions elicited by peer victimization (Khantzian, 1997). Longitudinal studies consistently support this meditational pathway in adolescents (Earnshaw et al., 2017; Hong et al., 2014; Marschall-lévesque et al., 2017; Meisel, Colder, Bowler, & Hussong, 2018; Rowe et al., 2019; Vannucci, Fagle, Simpson, & Ohannessian, 2020; Zapolski, Rowe, Fisher, Hensel, & Barnes-najor, 2018). For instance, structural equation modelling findings demonstrated that more frequent experiences of peer victimization in the 5th grade were associated with greater depressive symptoms in the 7th grade, which, in turn, were associated with a greater likelihood of alcohol use in the 10th grade (Earnshaw et al., 2017). In addition, Rowe et al. (2019) found a significant mediation path between 10th grade peer victimization, 11th grade anxiety symptoms, and 12^th^ grade alcohol use (similar results were found for the 9 to 11^th^ grade path).

Although most studies are based on the self-medication hypothesis, there is evidence of a temporal association between peer victimization and externalizing problems (Schoeler et al., 2018; Singham et al., 2017; van Lier et al., 2012). Schoeler et al (2018) found evidence of “*causal adverse effects*” on both internalizing and externalizing domains after taking these shared genetic influences into account. Similarly, there is well-validated literature on the effect of externalizing problems on alcohol use (Colder et al., 2018; Farmer et al., 2016; Steele et al., 1995). An alternative theory based on self-regulation hypotheses (the externalizing pathway), posits that the instability caused by peer victimization may generate hostile social-cognitive biases (hostile automatic thoughts, dysfunctional inhibitory control) that reinforces the development of externalizing behaviors, which further engage peer-victimized adolescents more frequently in alcohol use than other adolescents (Dodge et al., 1990). To our knowledge, the externalizing path between peer victimization and alcohol use has only been tested in one prospective study. Contrary to expectations, there was a significant association between peer victimization at age 14 and externalizing symptoms at age 18, but externalizing symptoms at age 18 did not significantly predict any further risk for alcohol misuse one year later (even after controlling for gender, age, family income and prior waves of alcohol use, externalizing, and internalizing symptoms). This suggests only concurrent relationships between externalizing and alcohol misuse symptoms (Meisel et al., 2018). When removing alcohol use at age 18 from the models, externalizing symptoms at age 18 significantly predict alcohol use at age 19. Thus, Meisel et al. (2018) failed to reveal an externalizing mediation path because of strong contemporaneous correlations (strong within time correlation at age 18), and possibly because the onset of substance use at earlier stages of adolescence was not captured by the study. Further investigations are needed. Thus, the current study hypothesized that peer victimization leads to alcohol use through two distinct time-varying mechanisms, the internalizing and externalizing pathways, by using a more sensitive developmental design allowing for the evaluation of temporal precedence in the onset of new victimization experiences and adolescent alcohol use.

### Personality as a moderator

The mechanisms of action linking peer victimization with alcohol use are certainly multifactorial and likely to differ across individual differences, reflecting pre-existing common vulnerabilities (Kavish et al., 2019). The possibility that individual differences exist in the strength of the associations has not been systematically explored, yet, it might have important implications for intervention. Research focusing on factors that moderate consequences of peer victimization is critical for understanding generalizability of a research finding (MacKinnon, 2011) and for supporting causal theories. It has been proposed that a number of potential moderators (e.g., gender, social support, attachment to school) could strengthen or weaken the association between peer victimization and substance use (Hong et al., 2014). Gender is the most studied moderator in this field of research that has concluded gender-related biases towards internalizing versus externalizing behaviors following victimization experiences; however, many studies on peer victimization have reported gender invariance (Forbes et al., 2019, 2020; Rosen et al., 2013; Schaefer et al., 2017). A growing body of research suggests that it may be less useful to take a gender-specific approach (Forbes et al., 2020), and so, other individual characteristics should be more investigated.

Personality might be a better choice as moderator, which have some gender variance (Castonguay-Jolin et al., 2013; Woicik et al., 2009), but might more directly related to psychopathology outcomes of victimization than gender (Afzali, Sunderland, et al., 2018; Calvete et al., 2016; Chinneck et al., 2018; Kaiser, Davis, Milich, & Smith, 2019; Kelly et al., 2018; Tani, Greenman, Schneider, & Fregoso, 1999). Due to the complexity and computational burden of moderated mediation models, few studies have explored the moderating role of personality from a developmental perspective. Those with sufficient sample sizes and developmental data (Brendgen et al., 2005; Calvete et al., 2016; Gallardo-Pujol & Pereda, 2013; Sugimura & Rudolph, 2012b; Zhu et al., 2016) have used multilevel analyses to reveal a weaker association between peer victimization and social anxiety for adolescents with high extraversion, fewer depressive symptoms in adolescent victims with low extraversion (Calvete et al., 2016). Another study reported a larger effect of peer victimization on behavioral problems among adolescents with high impulsivity compared to adolescents with low impulsivity (Zhu et al., 2016). These findings remain limited by the lack of a unified framework, and the limited set of personality tested, thus limiting the ability to confirm specificity of effects and potential personality-specific pathways between peer victimization and alcohol misuse. The current study aims to fill the third gap of the literature; hypothesising that peer victimization leads to alcohol use under the conditional influence of personality profiles.

### The current study

The current study provides a unique opportunity to apply a multilevel moderated mediation (conditional indirect effect) analysis to examining the specificity (personality), of multiple pathways (internalizing, externalizing paths) from peer victimization to alcohol use and misuse. Despite the fact that moderated mediation models can extract more information than simplistic models, few studies ventured into these models, in part due to lack of large longitudinal dataset and because of the difficulty of specifying and interpreting these models (MacKinnon, 2011). Yet, moderated mediation analysis can help quantify more complicated hypotheses, force consideration of alternative interpretations of the results, and lead to better research designs and more information gleaned from the study (Preacher, 2015). Moreover, the design of the present study is rather unique, first, because of the large population-based sample of 3,800 adolescents followed over 5 years, and second, because of the advanced computational method, the Bayesian multilevel method, that can establish temporal precedence between variables. In the absence of the possibility to use experimental designs to establish time-varying associations of peer victimization and alcohol use, developmental psychopathologists have now turned to the use of new computational methods in the form of multilevel models (MLMs), to establish temporal precedence in the relationship between two variables. Such models provide a rigorous test of predominance between two outcomes by quantifying the temporal association over multiple follow-up periods and by differentiating between-person, within-person, and lagged-within-person variance. This study investigated these effects while examining the “how” (mediation effect) and the “when” (moderation effect). In that respect, this is a rather markedly informative design enabling to test multiple concepts into one single model.

The current study examines broad levels of victimization, although many studies focus on specific form of victimization (i.e., physical, relational, and verbal) and its association with mental health. Results from a very well conducted study indicated that physical, verbal, and relational victimization had similar strength in associations across all levels of hierarchical model of psychopathology: specific diagnostics, internalizing versus externalizing dimensions, general factor of psychopathology (Forbes et al., 2020). Therefore, broad level of victimization are chosen in this study. Similarly, the use of dimensional structure of psychopathology (internalizing and externalizing spectra) instead of discrete diagnostic categories (e.g., depression, anxiety, conduct, and hyperactivity problems) was carefully selected based on evidence of the well-validated internalizing-externalizing model (Conway et al., 2019; Forbes et al., 2016). When Forbes et al. (2020) investigated the robustness of associations between levels of hierarchical model, the strongest relationships were at the level of the transdiagnostic internalizing, externalizing, and general psychopathology factors compared to specific diagnostic. Thus, the internalizing and externalizing dimensions act as powerful pathway variables that might channel the effect of peer victimization to alcohol use. In the present study, the internalizing factor represents the overlap between depressive symptoms and anxiety; and the externalizing factor represents the overlap between conduct, and hyperactivity problems. Finally, the present study integrates a large set of four personality traits (anxiety sensitivity, hopelessness, impulsivity, sensation seeking,) as moderators. These personality dimensions are high-risk predictors of substance use and specific psychopathology outcomes (Castellanos-Ryan, O’Leary-Barrett, Sully, & Conrod, 2013). Thus, these personality traits are good candidates for conditional factors of the association between peer victimization and alcohol use.

To conclude, the current study has three goals: (1) to investigate time-varying relationships between peer victimization and alcohol use; (2) through internalizing and externalizing pathways; (3) conditional to the levels of personality profiles of adolescents over a 5-year period.

## Methods

### Participants

Data from an ongoing population-based randomized controlled trial (Co-Venture) trial investigating the effectiveness of a 5-year personality-targeted drug and alcohol prevention program were used (O’Leary-Barrett et al., 2017). A large sample of adolescents (n = 3,800, 49.2% female, mean age = 12.8, SD = 0.4 years) was recruited from 31 schools in the Greater Montreal area. This sample of adolescents studied annually from 7th grade through 11th grade and is epidemiologically representative of each of its school districts regarding average size and socioeconomic index. The sample of schools represents 15% of all schools across the greater Montreal area and each of their respective school districts in size and deprivation indexes within 1.5 standard deviations. Two exclusion criteria for schools were specified: the school had to agree to the study protocol and the school could not have more than 50% of its seventh-grade students having special educational needs. Most of the data collection took place in the spring each year.

Participants were invited to complete a confidential annual web-based survey during class time intended to assess clinical, cognitive and behavioral information. Confidentiality was accomplished by emphasizing that parents and teachers would not have access to the survey results and by automatically anonymizing the assessments. Among the 3,800 adolescents who were invited to complete the survey annually, 3,612 (94.4%) who passed the quality control of the different questionnaires and provided consistent minimal demographic information(sex, age, SES) were included in the analysis. Participants who provided odd response patterns (e.g., same response for every question) or unusually fast reaction time (i.e., having a mean RT of 200 ms throughout the task, 5.6%) were excluded from the study. Excluded participants were not significantly different from others in demographic information (sex, age, SES). Ethical approval was obtained from the ethical review board at the [blinded]. All participants were included in the final study sample and analyses if 75% of their data across all items and annual survey waves were complete and reliable. Because more than 80 % of the participants did not receive the targeted personality-based intervention, which was a non-intrusive 2 x 90 minutes workshops, all participants were included in the final study sample. A sensitivity analysis will be conducted when approval to isolate data from non intervention schools will be granted. Attrition during the 5-year follow-up was 32% and was not significantly associated with participants’ characteristics.

### Measures

#### Independent variables

##### Peer victimization

Peer victimization was measured by asking the participants to retrospectively report their experiences in the past 12 months using the validated and widely used Olweus Peer/Victim Questionnaire—BVQ (Lee & Cornell, 2009). This questionnaire includes 4 questions on victimization (e.g., “I was called names, was made fun of, or teased in a hurtful way”). Participants were asked to rate their response on a 5-point Likert scale (0 = never, 1 = only once or twice, 2 = two or three times a month, 3 = once a week, 5 = several times a week).

##### Personality risk profiles

Four personality traits were assessed using the Substance Use Risk Profile Scale (Woicik et al., 2009). The SURPS is a 23-item questionnaire measuring personality risk for substance use and other behavioural problems according to four traits: anxiety– sensitivity (AS), described as a fear of anxiety-related physical sensations (e.g., “I get scared when I’m too nervous”), hopelessness (HOPE), a tendency towards low mood, worthlessness and negative beliefs about oneself, the world and the future (e.g., “I feel that I’m a failure”), sensation seeking (SS), defined by a low tolerance to boredom, a strong need for stimulation, and a willingness to take risks for the sake of having novel and varied experiences (e.g., “I enjoy new and exciting experiences even if they are unconventional”); and impulsivity (IMP), characterized by unplanned responses to internal or external stimulation or fast responses to given stimuli without deliberation and evaluation of consequences (e.g., “I often don’t think through before I speak”) (Newton et al., 2016). This instrument has good concurrent, predictive and incremental validity with regards to differentiating individuals prone to reinforcement-specific patterns of substance use and has been shown to be sensitive and specific with respect to predicting future substance misuse and other mental health problems in adolescents (Castellanos-Ryan et al., 2013; Krank et al., 2011). Each personality trait was assessed using 5–7 items each rated on a 4-point scale (1 = strongly disagree, 2 = disagree, 3 = agree, 4 = strongly agree) (Castonguay-Jolin et al., 2013). Reliability alpha coefficients and score ranges for each subscale were as follows, anxiety sensitivity (range = 5-20, α = 0.76), hopelessness (range = 7-28, α = 0.85), impulsivity (rang e= 5-20, α = 0.79), and sensation seeking (range = 6-24, α = 0.73).

#### Mediators

##### Externalizing problems

Externalizing symptoms were measured by asking participants to rate the occurrence of conduct and hyperactivity problems within the last 12 months using the Strengths and Difficulties Questionnaire (Goodman, 2001).The SDQ is one of the most commonly used instruments for screening psychopathology in children and adolescents. SDQ has been widely validated in various community and clinical samples across different countries (Giannakopoulos et al., 2013). Each domain was assessed using five self-reported items rated on a 3-point scale, such as “I get very angry and often lose my temper” for conduct problems, and “I am easily distracted, I find it difficult to concentrate” for hyperactivity problems (0 = not true, 1 = somewhat true, 2 = certainly true). Reliability alpha coefficients for each subscale were as follows, conduct problems (range = 0-10, α = 0.62), hyperactivity (range = 0-10, α = 0.73). The externalizing problems variable was created using the sum of the total scores of conduct and hyperactivity problems.

##### Internalizing symptoms

Internalizing symptoms were measured by asking participants to indicate to what extend they experienced depressive and anxiety symptoms over the past 12 months using the Brief Symptoms Inventory. Subscales consists of 7 self-reported items of depressive symptoms (e.g., “Feeling no interest in things”) and five self-reported items of anxiety symptoms (e.g., “Spells of terror or panic”) rated on a 5-point scale (0 = not at all, 1 = a little bit, 2 = moderately, 3 = quite a bit, 4 = often). Reliability alpha coefficients for each subscale were as follows, depression (range = 0-28, α = 0.88), anxiety (range = 0-20, α = 0.87). The internalizing symptoms variable was created using the sum of the total scores of depressive and anxiety symptoms. Although the SDQ covers an emotional symptoms scale, it does not discriminate depressive, phobic, anxiety, and obsessive-compulsive disorders unlike the BSI, that is also more widely used to identify clinically relevant psychological symptoms.

#### Dependent variable

##### Alcohol use

Alcohol use was assessed using the validated ‘Detection of Alcohol and Drug Problems in Adolescents’ questionnaire (Landry et al., 2004). The DEP-ADO reliably identifies youth with alcohol and drug use disorders (onset, frequency, quantity and harm associated with alcohol and drug use) in the past 12 months. Scores are summed, and a substance use disorder is identified when total scores are greater than 20. The DEP-ADO has demonstrated good construct validity, internal consistency, test retest and intermodel execution reliability in Quebec youth (Landry et al., 2004). Self-report measures have been found to have excellent discrimination (Clark & Winters, 2002) and predictive validity (White & Labouvie, 1989) with regards to adolescent substance use and problems (Castellanos & Conrod, 2006; Conrod, Castellanos-Ryan, & Strang, 2010; Conrod, Castellanos, & Mackie, 2008; Conrod et al., 2006). Participants rated their frequency of use on a 6-point scale (0 = Never, 1 = Occasionally, 2 = Monthly, 3 = 2–3 times per month, 4 = Weekly, 5 = Every day).

#### Covariates

Each model controlled for baseline socio-economic status (SES) and gender (0 = female, 1 = male). SES was assessed using the Family Affluence Scale for adolescents (Currie, Elton, Todd, & Platt, 1997).

### Analytical strategy

Data was analyzed using *Mplus 8.4* statistical software. Bayesian multilevel modelling was conducted to assess conditional indirect effects (i.e., moderated mediation analysis) with peer victimization as the independent variable, psychopathology as mediators, personality risk profiles as moderators, and alcohol use as the dependent variable (Figure 1). It has been demonstrated that performance of Bayesian methods yield higher power (e.g., unbiased estimates) compared to other traditional frequentist methods (e.g., maximum likelihood) to examine moderated mediation effects (Wang, Preacher, Wang, & Preacher, 2015).

**Figure 1.**
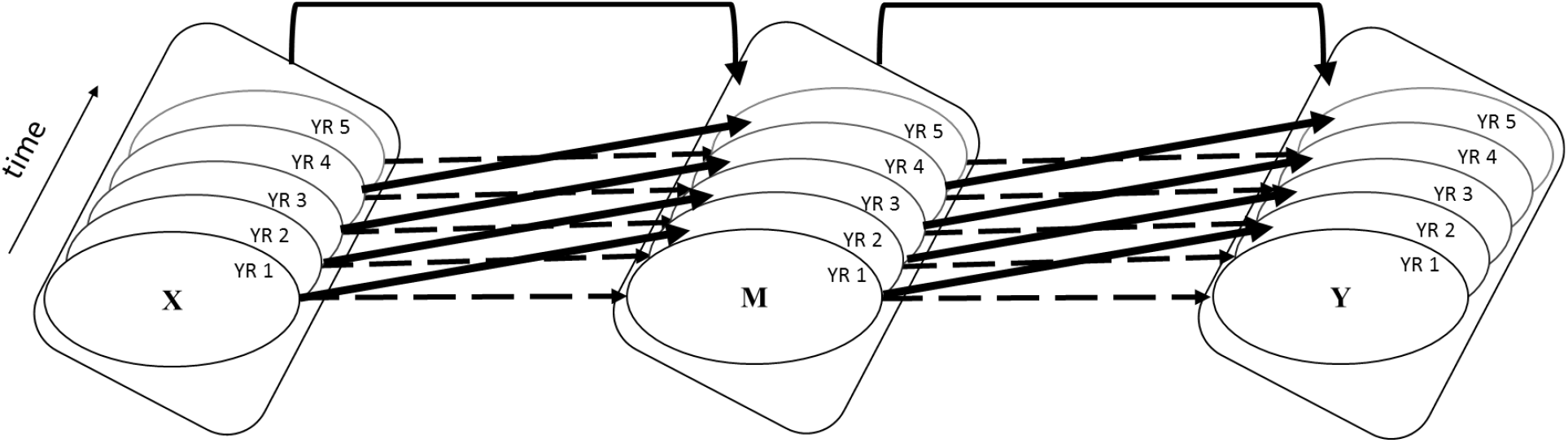
Multilevel Bayesian model assessing the indirect association of peer victimization and alcohol use through psychopathology and the potential mediating role of personality risk profiles. **Note.** Time = the survey waves (YR = year). Curved arrows = between-person effects. Dashed arrows = within-person concurrent effects. Diagonal arrows = lagged-within-person effects. (Model 1a: X = peer victimization, M = internalizing symptoms, Y= alcohol use; Model 1b: X = peer victimization, M = externalizing problems, Y = alcohol use; Model 2a: X = peer victimization high risk AS/HOPE, M = internalizing symptoms, Y = alcohol use; Model 2b: X = peer victimization high risk IMP/SS, M = externalizing problems, Y = alcohol use)

Moderated mediation models were conducted. Each model distinguished the role of three aspects of peer victimization: between-person effects (the effect of average peer victimization over 5 years), within-person concurrent effects (change in level of peer victimization within a given year compared to person’s mean peer victimization within that same year) and lagged-within-person effects (level of peer victimization the year before compared to person’s mean peer victimization the following year). The time parameter (i.e., year of assessment) was coded from one to five.

The data analytical approach consisted of two steps. The first step estimated the indirect association between the predictor (i.e., peer victimization) and the outcome variable (i.e., alcohol use) through each mediators (i.e., internalizing symptoms, externalizing problems) in two distinct models: model 1a assessed the indirect between-, within-, and lagged-within-person effects of peer victimization on alcohol use through internalizing symptoms; model 1b assessed the indirect between-, within-, and lagged-within-person effects of peer victimization on alcohol use through externalizing problems. The second step consisted of integrating all previously named parameters with the addition of the moderators (i.e., high AS/HOPE, high IMP/SS) in two distinct models: model 2a assessed the same parameters as in model 1a with the addition of high AS/HOPE as a moderator; model 2b assessed the same parameters as in model 1b with the addition of high IMP/SS as a moderator. Likewise, to assess the moderating role of low personality profiles, a new set of models were estimated following the same logic: model 3a assessed the same parameters as in model 1a with the addition of low AS/HOPE as a moderator; model 3b assessed the same parameters as in model 1b with the addition of low IMP/SS as a moderator.

The “model constraint” command was used in *Mplus* to calculate the indirect effects based on the product of component path coefficients. Standard errors and 95% credibility intervals for indirect effects were calculated. To increase the clarity of our results the means, standard deviation, and correlations between the main variables is presented in Table 1.

**Table 1.**
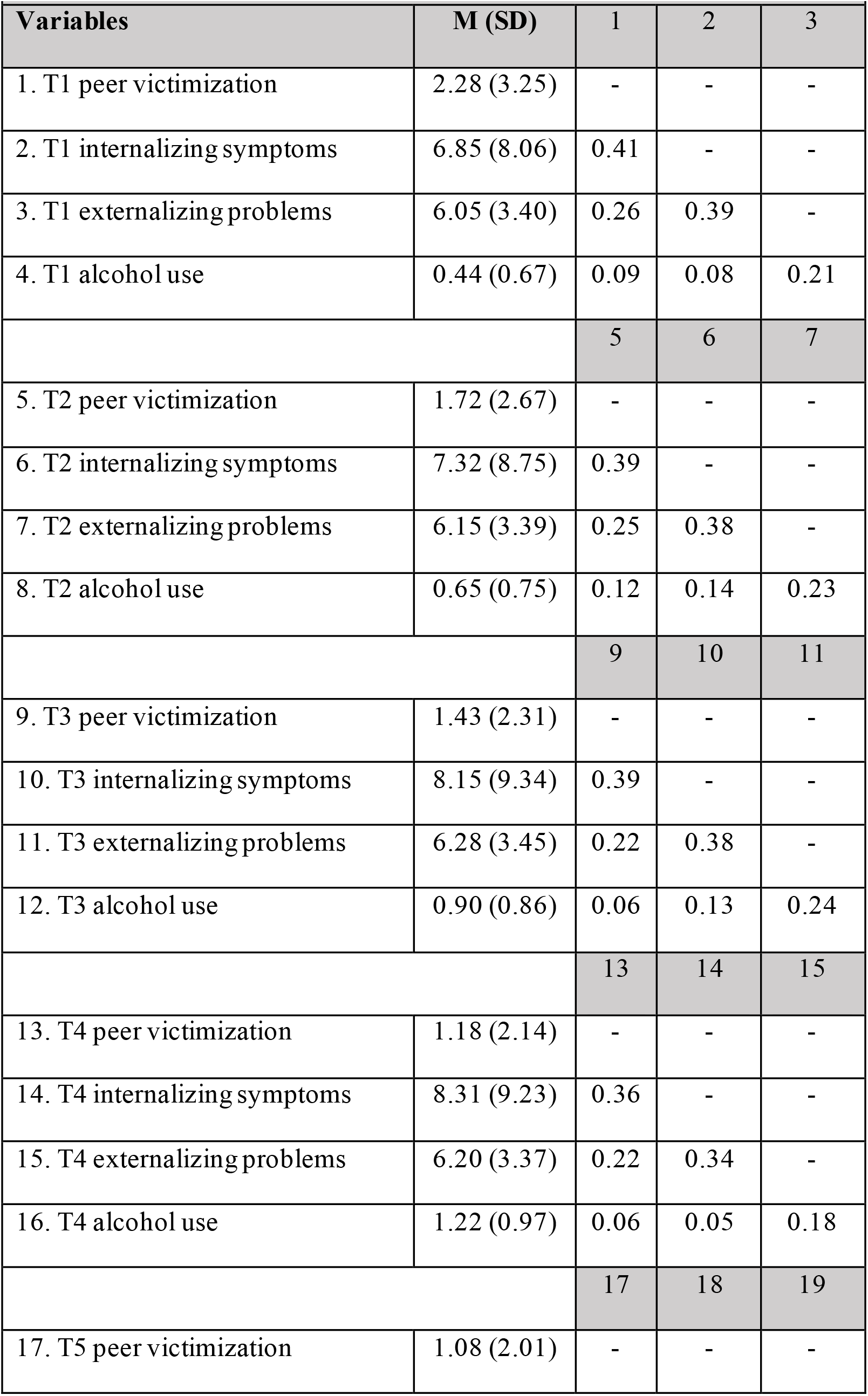
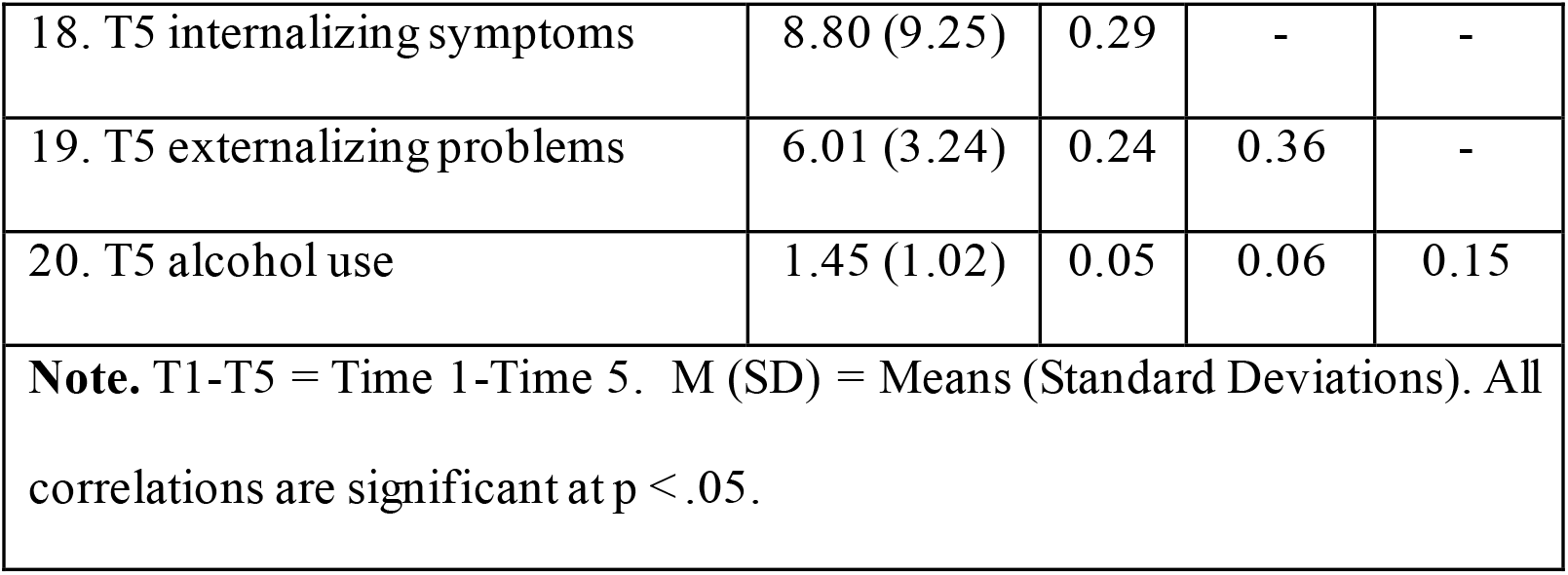
Means, standard deviations, and correlations among variables

| **Table 1. Means, standard deviations, and correlations among variables** |  |  |  |  |
| --- | --- | --- | --- | --- |
| **Variables** | **M (SD)** | **1** | **2** | **3** |
| 1. T1 peer victimization | 2.28 (3.25) | - | - | - |
| 2. T1 internalizing symptoms | 6.85 (8.06) | 0.41 | - | - |
| 3. T1 externalizing problems | 6.05 (3.40) | 0.26 | 0.39 | - |
| 4. T1 alcohol use | 0.44 (0.67) | 0.09 | 0.08 | 0.21 |
|  |  | **5** | **6** | **7** |
| 5. T2 peer victimization | 1.72 (2.67) | - | - | - |
| 6. T2 internalizing symptoms | 7.32 (8.75) | 0.39 | - | - |
| 7. T2 externalizing problems | 6.15 (3.39) | 0.25 | 0.38 | - |
| 8. T2 alcohol use | 0.65 (0.75) | 0.12 | 0.14 | 0.23 |
|  |  | **9** | **10** | **11** |
| 9. T3 peer victimization | 1.43 (2.31) | - | - | - |
| 10. T3 internalizing symptoms | 8.15 (9.34) | 0.39 | - | - |
| 11. T3 externalizing problems | 6.28 (3.45) | 0.22 | 0.38 | - |
| 12. T3 alcohol use | 0.90 (0.86) | 0.06 | 0.13 | 0.24 |
|  |  | **13** | **14** | **15** |
| 13. T4 peer victimization | 1.18 (2.14) | - | - | - |
| 14. T4 internalizing symptoms | 8.31 (9.23) | 0.36 | - | - |
| 15. T4 externalizing problems | 6.20 (3.37) | 0.22 | 0.34 | - |
| 16. T4 alcohol use | 1.22 (0.97) | 0.06 | 0.05 | 0.18 |
|  |  | **17** | **18** | **19** |
| 17. T5 peer victimization | 1.08 (2.01) | - | - | - |
| 18. T5 internalizing symptoms | 8.80 (9.25) | 0.29 | - | - |
| 19. T5 externalizing problems | 6.01 (3.24) | 0.24 | 0.36 | - |
| 20. T5 alcohol use | 1.45 (1.02) | 0.05 | 0.06 | 0.15 |
| **Note.** T1-T5 = Time 1-Time 5. M (SD) = Means (Standard Deviations). All correlations are significant at $p < .05$. | | | | |

## Results

### The indirect association of peer victimization and alcohol use through internalizing symptoms (Model 1a)

The results (Table 2) indicated significant between-person mediation effect of peer victimization (*p* < .0001) on alcohol use through internalizing symptoms over the five-year period, independent of within effects, SES, and gender: on average over the 5-year period, those prone to higher levels of peer victimization are also prone to higher levels of alcohol and use, and this relationship was mediated by high overall levels of internalizing symptoms. Over and above the significant between effect, results also indicated a significant within-person mediation effect (*p* < .0001) and lagged-within-person mediation effect (*p* < .01): any further increases in exposure to peer victimization in a given year was associated with increased risk of developing internalizing symptoms, and subsequently, alcohol use during the same year and one year later.

**Table 2.** Model 1a: Estimated parameters for multilevel models assessing internalizing symptoms as mediator of the temporal association between victimization and alcohol use

| Predictors | Estimate | St. Er. | Pr(>\|t\|) | 95% CI |
| --- | --- | --- | --- | --- |
| Intercept (INT) | 6.133 | 0.373 | 0.000 | 5.387, 6.859 |
| Intercept (ALC) | -0.394 | 0.043 | 0.000 | -0.477, -<br>0.309 |
| Time (INT) | 0.738 | 0.035 | 0.000 | 0.674, 0.809 |
| Time (ALC) | 0.264 | 0.004 | 0.000 | 0.257, 0.271 |
| Gender | 0.121 | 0.025 | 0.000 | 0.070, 0.171 |
| SES | 0.069 | 0.006 | 0.000 | 0.057, 0.081 |
| Victimization between on ALC | 0.026 | 0.010 | **0.005** | 0.007, 0.045 |
| Victimization within on ALC | 0.010 | 0.003 | **0.002** | 0.003, 0.016 |
| Victimization lagged on ALC | -0.004 | 0.004 | 0.191 | -0.011, 0.004 |
| Victimization between on INT | 1.870 | 0.076 | **0.000** | 1.721, 2.013 |
| Victimization within on INT | 0.806 | 0.033 | **0.000** | 0.742, 0.871 |
| Victimization lagged on INT | 0.185 | 0.038 | **0.000** | 0.111, 0.259 |
| INT between on ALC | 0.011 | 0.003 | **0.000** | 0.006, 0.015 |
| INT within on ALC | 0.006 | 0.001 | **0.000** | 0.004, 0.008 |
| INT lagged on ALC | 0.003 | 0.001 | **0.014** | 0.000, 0.005 |

**Note.** Significant effects are marked in **bold** font (one-tailed p-value). SES = socio-economic status. INT = internalizing symptoms. ALC = alcohol use. Estimates were calculated using unstandardized beta.

### The indirect association of peer victimization and alcohol use through externalizing problems (Model 1b)

Regarding indirect associations of peer victimization on alcohol use through externalizing problems, the results are presented in Table 3. There was a significant between-person mediation effect of peer victimization (*p* < .0001), on alcohol use, over and above within effects, SES, and gender. The results also indicated significant within-person mediation effect (*p* < .0001) and lagged-within-person mediation effect(*p* < .0001), while controlling for the between-person effect.

**Table 3.** Model 1b: Estimated parameters for multilevel models assessing externalizing problems as mediator of the temporal association between victimization and alcohol use

| Predictors | Estimate | St. Er. | Pr(>\|t\|) | 95% CI |
| --- | --- | --- | --- | --- |
| Intercept (EXT) | 4.752 | 0.168 | 0.000 | 4.425, 5.076 |
| Intercept (ALC) | -0.622 | 0.046 | 0.000 | -0.712, -0.535 |
| Time (EXT) | 0.113 | 0.014 | 0.000 | 0.086, 0.141 |
| Time (ALC) | 0.266 | 0.004 | 0.000 | 0.258, 0.274 |
| Gender | 0.062 | 0.021 | 0.003 | 0.021, 0.105 |
| SES | 0.065 | 0.006 | 0.000 | 0.053, 0.077 |
| Victimization between on ALC | 0.010 | 0.009 | 0.128 | -0.007, 0.0227 |
| Victimization within on ALC | 0.011 | 0.003 | **0.002** | 0.004, 0.018 |
| Victimization lagged on ALC | -0.004 | 0.004 | 0.189 | -0.011, 0.004 |
| Victimization between on EXT | 0.575 | 0.036 | **0.000** | 0.504, 0.643 |
| Victimization within on EXT | 0.156 | 0.012 | **0.000** | 0.132, 0.179 |
| Victimization lagged on EXT | 0.075 | 0.015 | **0.000** | 0.044, 0.103 |
| EXT between on ALC | 0.061 | 0.005 | **0.000** | 0.052, 0.071 |
| EXT within on ALC | 0.028 | 0.003 | **0.000** | 0.023, 0.034 |
| EXT lagged on ALC | 0.016 | 0.003 | **0.000** | 0.009, 0.023 |

**Note.** Significant effects are marked in **bold** font. SES= socio-economic status. EXT= externalizing problems. ALC = alcohol use. Estimates were calculated using unstandardized beta.

### The moderating effect of anxiety sensitivity and hopelessness between peer victimization and alcohol use (Models 2a, 3a)

Results are presented in Table 4, model 2a show a significant conditional indirect between-person effect (*p* < .0001) and within-person effect (*p* < .0001) of peer victimization on alcohol use through internalizing symptoms for adolescents high in anxiety sensitivity or hopelessness. No conditional indirect lagged-within-person effect was found (*p* > .05). Conversely, the indirect between-person, within-person and lagged-within-person effects (*p* > .05) did not reach significance for those with low levels of these profiles. Thus, the indirect relationship between peer victimization and alcohol through internalizing symptoms differs across levels of anxiety sensitivity and hopelessness.

**Table 4.**
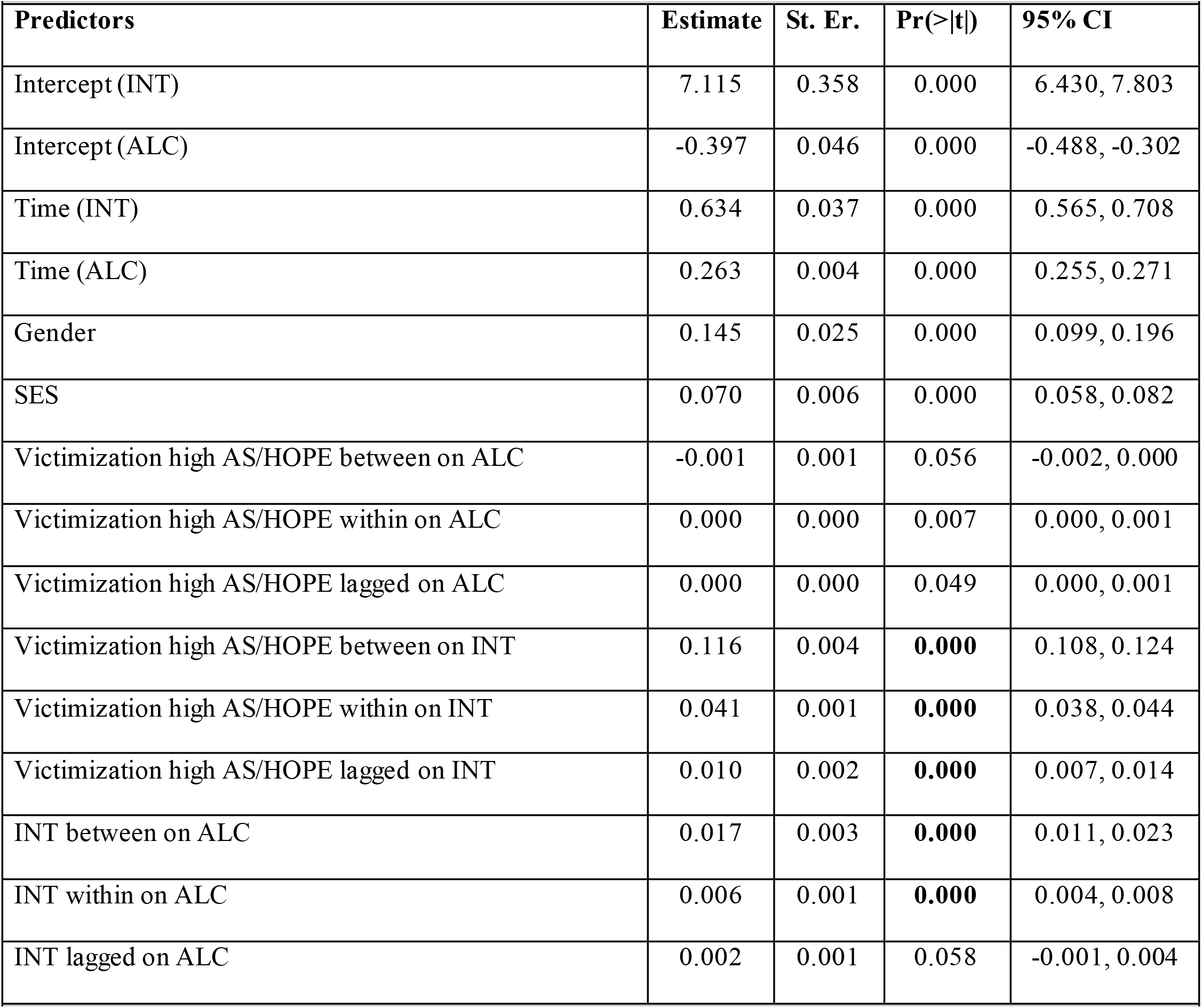
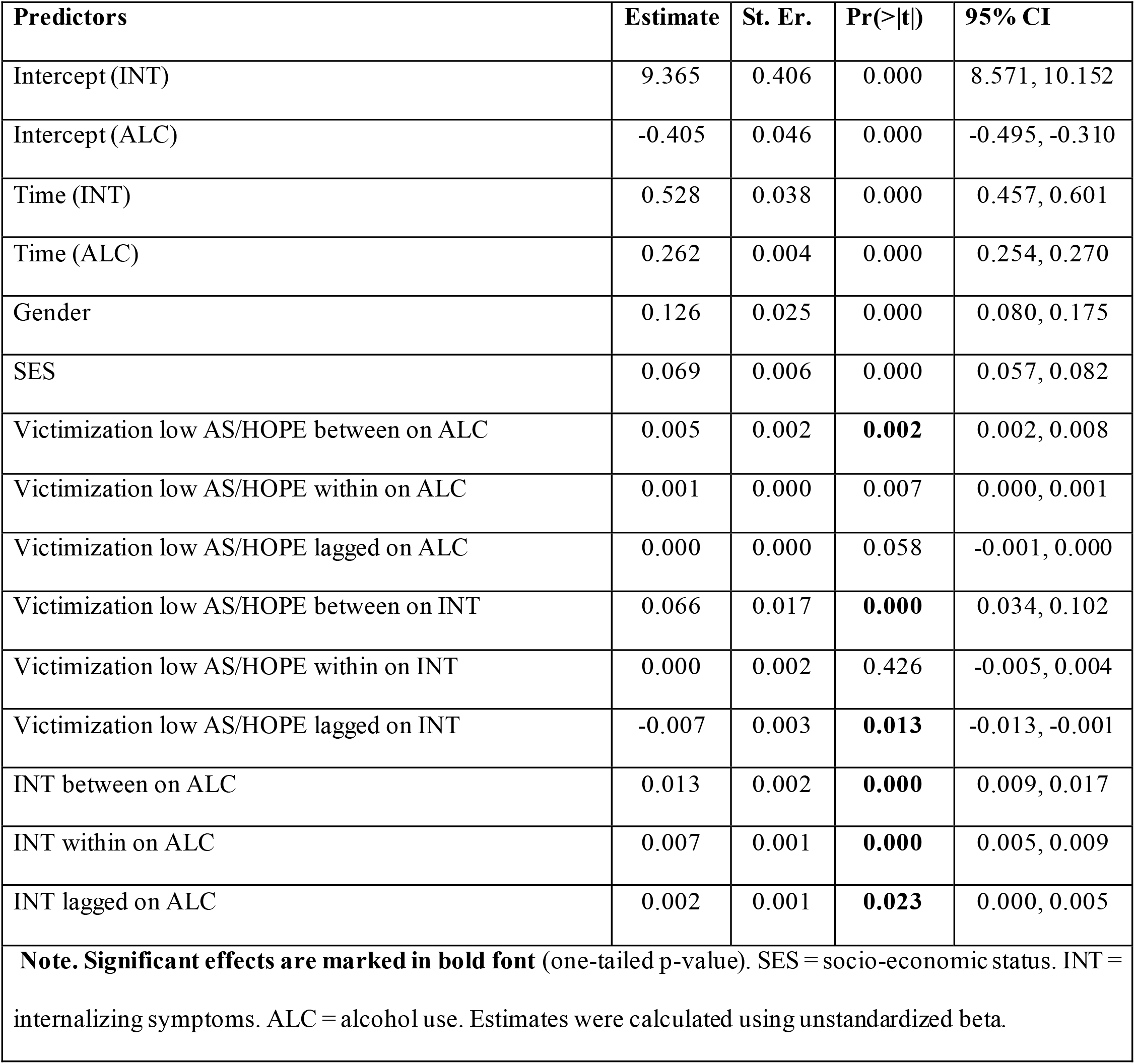
Model 2a Estimated parameters for multilevel models assessing internalizing symptoms as mediator of the temporal association between victimization with high risk profile (anxiety sensitivity and hopelessness) and alcohol use Model 3a Estimated parameters for multilevel models assessing internalizing symptoms as mediator of the temporal association between victimization with low risk profile (anxiety sensitivity and hopelessness) and alcohol use

### The moderating effect of impulsivity and sensation seeking between peer victimization and alcohol use through externalizing problems (Models 2b, 3b)

Results (Table 5) from the model 2b indicated that there was a conditional indirect effect of peer victimization on alcohol use through externalizing problems : between-person, within-person, and lagged-within-person effects (*p* < .0001). Conversely, results from the model 3b showed no significant indirect between-person, within-, and lagged-within-person (*p* >.05) effects for adolescents low on these profiles. In other words, the effect of peer victimization on alcohol use through externalizing problems is conditioned for adolescents with high level of impulsivity and sensation seeking. Thus, the indirect relationship between peer victimization and alcohol through externalizing problems differs across levels of impulsivity and sensation seeking.

**Table 5.** Model 2b Estimated parameters for multilevel models assessing externalizing symptoms as mediator of the temporal association between victimization with high risk profile (impulsivity and sensation seeking) and alcohol use Model 3b Estimated parameters for multilevel models assessing externalizing symptoms as mediator of the temporal association between victimization with low profile (impulsivity and sensation seeking) and alcohol use

| **Table 5.** |  |  |  |  |
| --- | --- | --- | --- | --- |
| **Model 2b Estimated parameters for multilevel models assessing externalizing symptoms as mediator of the temporal association between victimization with high risk profile (impulsivity and sensation seeking) and alcohol use** |  |  |  |  |
| **Predictors** | **Estimate** | **St. Er.** | **Pr(>\|t\|)** | **95% CI** |
| Intercept (EXT) | 5.206 | 0.160 | 0.000 | 4.911, 5.523 |
| Intercept (ALC) | -0.578 | 0.047 | 0.000 | -0.668, -0.485 |
| Time (EXT) | 0.090 | 0.012 | 0.000 | 0.066, 0.114 |
| Time (ALC) | 0.265 | 0.004 | 0.000 | 0.258, 0.272 |
| Gender | 0.062 | 0.020 | 0.002 | 0.021, 0.101 |
| SES | 0.064 | 0.006 | 0.000 | 0.052, 0.076 |
| Victimization high risk IMP/SS between on ALC | 0.002 | 0.000 | **0.000** | 0.001, 0.003 |
| Victimization high risk IMP/SS within on ALC | 0.001 | 0.000 | 0.000 | 0.000, 0.001 |
| Victimization high risk IMP/SS lagged on ALC | 0.000 | 0.000 | 0.319 | 0.000, 0.000 |
| Victimization high risk IMP/SS between on EXT | 0.033 | 0.002 | **0.000** | 0.030, 0.037 |
| Victimization high risk IMP/SS within on EXT | 0.008 | 0.000 | **0.000** | 0.007, 0.009 |
| Victimization high risk IMP/SS lagged on EXT | 0.003 | 0.001 | **0.000** | 0.002, 0.004 |
| EXT between on ALC | 0.050 | 0.005 | **0.000** | 0.040, 0.060 |
| EXT within on ALC | 0.027 | 0.003 | **0.000** | 0.022, 0.032 |
| EXT lagged on ALC | 0.016 | 0.003 | **0.000** | 0.010, 0.022 |

| **Model 3b Estimated parameters for multilevel models assessing externalizing symptoms as mediator of the temporal association between victimization with low profile (impulsivity and sensation seeking) and alcohol use** |  |  |  |  |
| --- | --- | --- | --- | --- |
| **Predictors** | **Estimate** | **St. Er.** | **Pr(>\|t\|)** | **95% CI** |
| Intercept (EXT) | 5.787 | 0.174 | 0.000 | 5.457, 6.137 |
| Intercept (ALC) | -0.558 | 0.046 | 0.000 | -0.647, -<br>0.467 |
| Time (EXT) | 0.066 | 0.012 | 0.000 | 0.041, 0.090 |
| Time (ALC) | 0.264 | 0.004 | 0.000 | 0.256, 0.270 |
| Gender | 0.049 | 0.021 | 0.007 | 0.008, 0.090 |
| SES | 0.065 | 0.006 | 0.000 | 0.052, 0.077 |
| Victimization low IMP/SS between on ALC | -0.006 | 0.001 | **0.000** | -0.008, -<br>0.004 |
| Victimization low IMP/SS within on ALC | 0.000 | 0.000 | 0.453 | 0.000, 0.000 |
| Victimization low IMP/SS lagged on ALC | 0.000 | 0.000 | 0.426 | -0.001, 0.000 |
| Victimization low IMP/SS between on EXT | 0.006 | 0.005 | 0.113 | -0.004, 0.016 |
| Victimization low IMP/SS within on EXT | -0.001 | 0.001 | 0.043 | -0.003, 0.000 |
| Victimization low IMP/SS lagged on EXT | 0.000 | 0.001 | 0.468 | -0.002, 0.002 |
| EXT between on ALC | 0.064 | 0.004 | **0.000** | 0.056, 0.072 |
| EXT within on ALC | 0.029 | 0.003 | **0.000** | 0.024, 0.035 |
| EXT lagged on ALC | 0.015 | 0.003 | **0.000** | 0.009, 0.022 |
| **Note. Significant effects are marked in bold font (one-tailed p-value). SES = socio-economic status. EXT = externalizing problems. ALC = alcohol use. Estimates were calculated using unstandardized beta.** |  |  |  |  |

## Discussion

By employing a rather advanced statistical modelling approach with a large population-based sample of 3,800 adolescents, the present study, investigated the mediating role of psychopathology in the time-varying associations between peer victimization and alcohol use and the moderating role of personality risk profiles in this indirect association.

This study provides strong evidence for the existence of two indirect paths from peer victimization to alcohol use through internalizing symptoms and externalizing problems. The two models (model 1a, 1b) both revealed between-person effects. That is, adolescents exposed to peer victimization increase their general tendency toward alcohol use through the development of internalizing symptoms and externalizing problems, while controlling the effect of each variable across within-levels during the 5-year period. Beyond this general tendency, within-, and lagged-within-person effects were observed, meaning that changes in level of exposure to peer victimization within a given year predispose adolescents to further alcohol use through the exacerbation of internalizing symptoms and externalizing problems in that same year and one year later. Besides demonstrating strong evidence for temporal mediation effects, these latter results highlighted three important aspects of these mediations: they are general, immediately experienced and longer-lasting. These results are in line with previous studies for the internalizing path (Earnshaw et al., 2017; Marschall-lévesque et al., 2017; Rowe et al., 2019; Vannucci et al., 2020; Zapolski et al., 2018), and are conflicting with the results of the first study that investigated the externalizing path (Meisel et al., 2018), which did not find support for negative peer experiences operating through externalizing problems on alcohol use, although their initial assumption was in favor of the existence of the externalizing path. It is possible that adolescents exposed to peer victimization learn to become more aggressive by being repeatedly reinforced for hitting or calling back onto their aggressors (Renouf et al., 2010). Alternatively, repeated peer victimization may generate a cognitive style that reinforces negative evaluations of the self and the future, and may lead to hypervigilance and a tendency to overestimate the level of threat (Espelage & Holt, 2001; Wang, 2011). Future research should investigate differential cognitive styles for both internalizing and externalizing pathways.

This study then explored if personality risk profiles might explain differential reactivity to peer victimization. When integrating high levels of anxiety sensitivity and hopelessness (model 2a), there was a between-, and a within-person conditional indirect effect of peer victimization on alcohol use through internalizing symptoms. No lagged-within-person conditional indirect effect was found. This suggest that the strength of the indirect relation between peer victimization and alcohol use depend on whether adolescents have high anxiety sensitivity and hopelessness. Similar conclusion can be drawn when investigating the conditional effect of high versus low levels of impulsivity and sensation seeking in the indirect association between peer victimization and alcohol use through externalizing problems. High impulsivity and sensation seeking increased the indirect effect of peer victimization at every level (between, within, and lagged); whereas no such effects were found with the low impulsivity and sensation seeking condition. The findings suggest that exposure to peer victimization is more powerful in shaping externalizing problems towards alcohol use when adolescent individual characteristics are impulsivity and sensation seeking. This is in line with numerous studies demonstrating that personality factors are implicated in the vulnerability to adolescent alcohol use (Conrod et al., 2008) and other psychopathological outcomes (Castellanos & Conrod, 2006; Krueger et al., 1996). In this case, individual differences approach, such as focusing on personality-specific aspects, could be a major advantage when selecting, targeting, and assisting high-risk youth before they have initiated alcohol use. This approach will result in more sensitivity with respect to identifying current and future substance users. Overall, the current results suggest that peer victimization may act as a stressor, generating specific manifestation of psychopathology sensitive to personality profiles, which makes peer victimization pernicious and difficult to tackle.

The current study is not without limitations. First, self-reported measure of peer victimization were collected in a classroom during school hours. Because of the sensitive nature of reporting peer victimization surrounded by peers, it might have been underreported. To overcome this problem, trained staff supervised data collection, and therefore allowed a confidential context. Second, although we accounted for important potential confounders, it is possible that other factors, such as individual (e.g., puberty), familial (e.g., parenting), or social (e.g., peer drinking) factors might affect the associations observed. Additional research is needed to shed light into these interactions. Third, cybervictimization was not investigated; yet, it is also associated with internalizing, externalizing problems (Fisher et al., 2016), and substance use (Kowalski, Giumetti, Schroeder, & Lattanner, 2014). Fourth, the present study investigated only the unidirectional path from peer victimization to alcohol use; although, evidence for the reverse direction exists (Maniglio, 2017). Future studies should address this issue with bidirectional effects, such as cross-lagged path model, but such designs should be sensitive to shorter time-interval when investigating lagged effects.

Despite these limitations, a major strength of the current study is the use of a large community sample size of 3,800 adolescents followed during 5 consecutive years, enhancing confidence in the generalizability of the results. In contrast to generalizability, another main strength is the unique methodological design that depicts an overall picture of how and when peer victimization exposure tend to be the most harmful. The statistical model used was able to test specificity of effect by highlighting the importance of identifying high risk groups, to then enable to tailor intervention.

Intervention programs have usually limited resources to accomplish their goals. Intervention programs will cost less and provide greater benefits if the critical ingredients of interventions can be identified. Findings suggest that anti-peer programs should adopt a different approach by removing ineffective components (Zych et al., 2017) and adding targeted personality-based strategies to reduce the emergence of various psychopathological outcomes and prevent alcohol abuse later in life. Adolescents with different psychopathological patterns of victimization cannot be addressed with a uniform ‘one size fits all’ approach. Our previous research have brought new evidence in reducing victimization, internalizing symptoms, externalizing problems, and substance use through a selective intervention based on the four personality dimensions used in this study (Conrod, Topper, O’Leary-Barrett, & Afzali, 2019; Kelly et al., 2019; O’Leary-Barrett et al., 2013). For example, impulsivity-based intervention reduce conduct problems, and anxiety sensitivity-based intervention reduce anxiety symptoms (O’Leary-Barrett et al., 2013). There was higher levels of victimization among adolescents identified by personality risk, and the magnitude of decrease in victimization was higher among students who participated in the intervention (Conrod et al., 2019). Moreover, receiving the personality-based intervention was beneficial for adolescents who experienced peer victimization regarding their alcohol-related harm compared to non-victimized adolescents (Edalati et al., 2019).

The current study proposes a novel developmentally informed model to push research beyond a focus on simple cross-sectional associations and specific diagnostic pathology. The findings of the current study stress the need to regulate peer behavior. Individual’s own characteristics (personality profiles) and environmental factors (peer victimization) appear to be a dangerous combination for overall, short-term, and long-term risks of developing psychopathology, and, further along engaging in alcohol use more severely than others who do not have these characteristics. In this study, new perspectives are provided by addressing the specifics how and for whom personality profiles can shape the immediate and long-term consequences of peer victimization over the course of adolescence.

